# BioKIT: a versatile toolkit for processing and analyzing diverse types of sequence data

**DOI:** 10.1101/2021.10.02.462868

**Authors:** Jacob L. Steenwyk, Thomas J. Buida, Carla Gonçalves, Dayna C. Goltz, Grace Morales, Matthew E. Mead, Abigail L. LaBella, Christina M. Chavez, Jonathan E. Schmitz, Maria Hadjifrangiskou, Yuanning Li, Antonis Rokas

## Abstract

Bioinformatic analysis—such as genome assembly quality assessment, alignment summary statistics, relative synonymous codon usage, paired-end aware quality trimming and filtering of sequencing reads, file format conversion, and processing and analysis—is integrated into diverse disciplines in the biological sciences. Several command-line pieces of software have been developed to conduct some of these individual analyses; however, the lack of a unified toolkit that conducts all these analyses can be a barrier in workflows. To address this obstacle, we introduce BioKIT, a versatile toolkit for the UNIX shell environment with 40 functions, several of which were community-sourced, that conduct routine and novel processing and analysis of genome assemblies, multiple sequence alignments, coding sequences, sequencing data, and more. To demonstrate the utility of BioKIT, we assessed the quality and characteristics of 901 eukaryotic genome assemblies, calculated alignment summary statistics for 10 phylogenomic data matrices, determined relative synonymous codon usage across 171 fungal genomes including those that use alternative genetic codes, and demonstrate that a novel metric, gene-wise relative synonymous codon usage, can accurately estimate gene-wise codon optimization. BioKIT will be helpful in facilitating and streamlining sequence analysis workflows. BioKIT is freely available under the MIT license from GitHub (https://github.com/JLSteenwyk/BioKIT), PyPi (https://pypi.org/project/jlsteenwykbiokit/), and the Anaconda Cloud (https://anaconda.org/jlsteenwyk/jlsteenwyk-biokit). Documentation, user tutorials, and instructions for requesting new features are available online (https://jlsteenwyk.com/BioKIT).

## Introduction

Bioinformatics is the application of computational tools to process and analyze biological data, such as nucleotide or amino acid sequences in the form of genome assemblies, gene annotations, and multiple sequence alignments (Bayat, 2002). Diverse disciplines in the biological sciences rely on bioinformatic methods and software (Wren, 2016). Recently, researchers have acknowledged the need to consider diverse types of biological scientists with different levels of experience when developing software (Kumar and Dudley, 2007). It is also essential to implement high standards of software development that ensure software functionality and archival stability (Mangul, Mosqueiro, *et al*., 2019; Mangul, Martin, *et al*., 2019). For example, code quality can be improved by utilizing unit and integration tests, which ensure faithful function of code (Darriba *et al*., 2018). As a result, the development of effective and user-friendly software for diverse biologists often requires an interdisciplinary team of software engineers, biologists, and others.

Even though numerous bioinformatic pieces of software are available, there are still several barriers to creating seamless and reproducible workflows (Kim *et al*., 2018). This issue in part stems from different pieces of software requiring different input file formats, being unable to account for non-standard biological phenomena such as the use of alternative genetic codes, or can only be executed using web servers or graphical user interfaces, which cannot be incorporated into high-throughput pipelines. Another factor is that multiple pieces of software or custom scripts are typically needed to execute different steps in a larger bioinformatic pipeline; for example, bioinformatic workflows often rely on one software/script for converting file formats, another software/script for translating sequences using standard and non-standard genetic codes, another software/script to examine the properties of genomes or multiple sequence alignments, and so on. As a result, maintaining efficacious bioinformatic workflows is cumbersome (Kulkarni *et al*., 2018). Thus, the bioinformatic community would benefit from a multi-purpose toolkit that contains diverse processing and analysis functions.

To address this need, we—an interdisciplinary team of software engineers, evolutionary biologists, molecular biologists, microbiologists, and others—developed BioKIT, a versatile toolkit with 40 functions, several of which were community sourced, that conduct routine and novel processing and analysis of diverse sequence files including genome assemblies, multiple sequence alignments, protein coding sequences, and sequencing data (Table 1). Functions implemented in BioKIT facilitate a wide variety of standard bioinformatic analyses, including genome assembly quality assessment (e.g., N50, L50, assembly size, guanine-cytosine (GC) content, number of scaffolds, and others), the calculation of multiple sequence alignment properties (i.e., number of taxa, alignment length, the number of constant sites, the number of parsimony-informative sites, and the number of variable sites), and processing and analysis of protein coding sequences (e.g., translation using 26 genetic codes including user-specified translation tables, GC content at the first, second, and third codon positions, and relative synonymous codon usage). To demonstrate the utility of BioKIT, we examined the genome assembly quality of 901 eukaryotic genomes, evaluated the properties of 10 phylogenomic data matrices, calculated relative synonymous codon usage in 171 fungal genomes, and estimated codon optimization in each gene from two *Saccharomyces* budding yeast species using a novel metric, gene-wise relative synonymous codon usage (gw-RSCU). BioKIT comes complete with common and novel functions that will help improve reproducibility and accessibility of diverse bioinformatic analysis and facilitate discovery in the biological sciences.

**Table 1.**
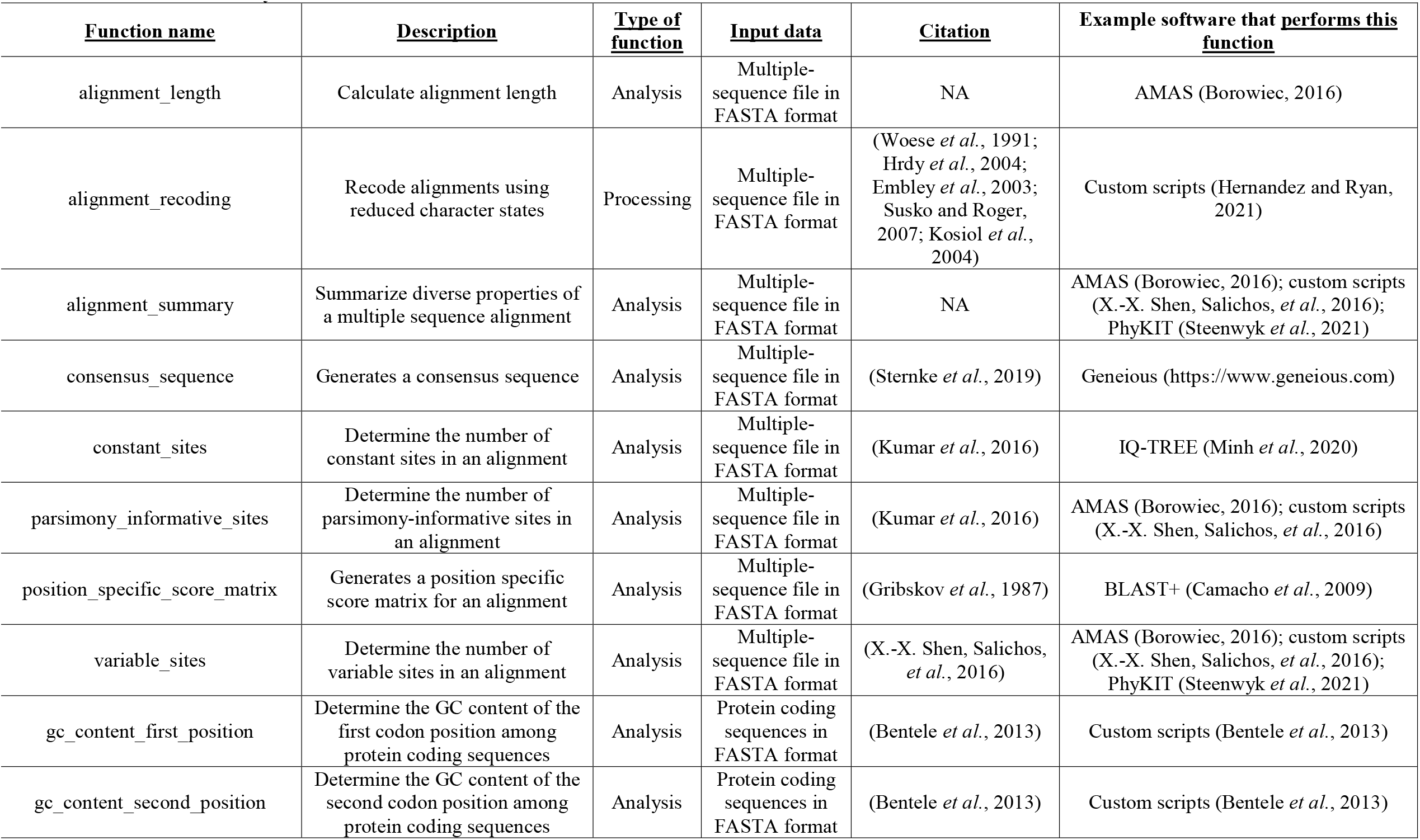
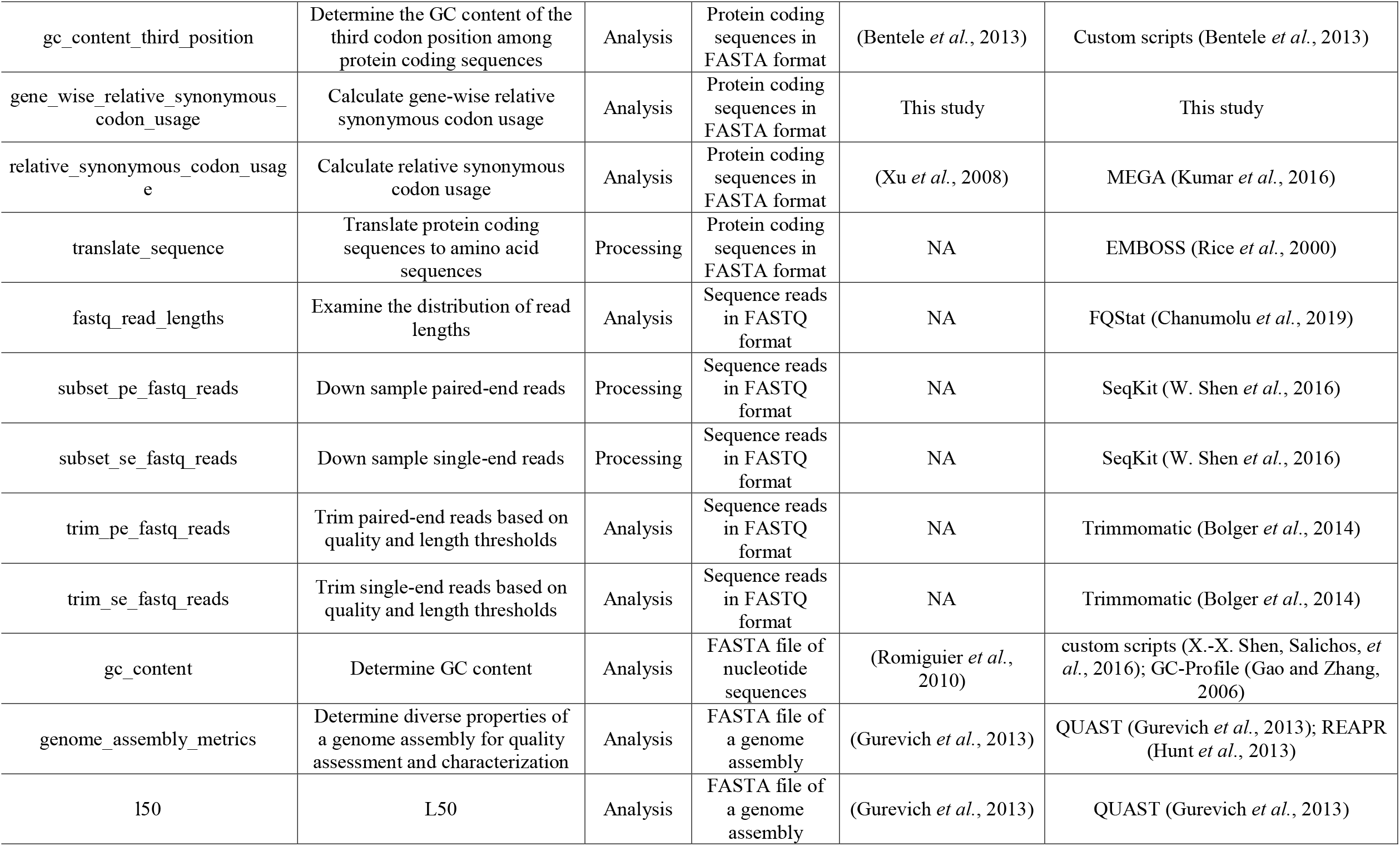
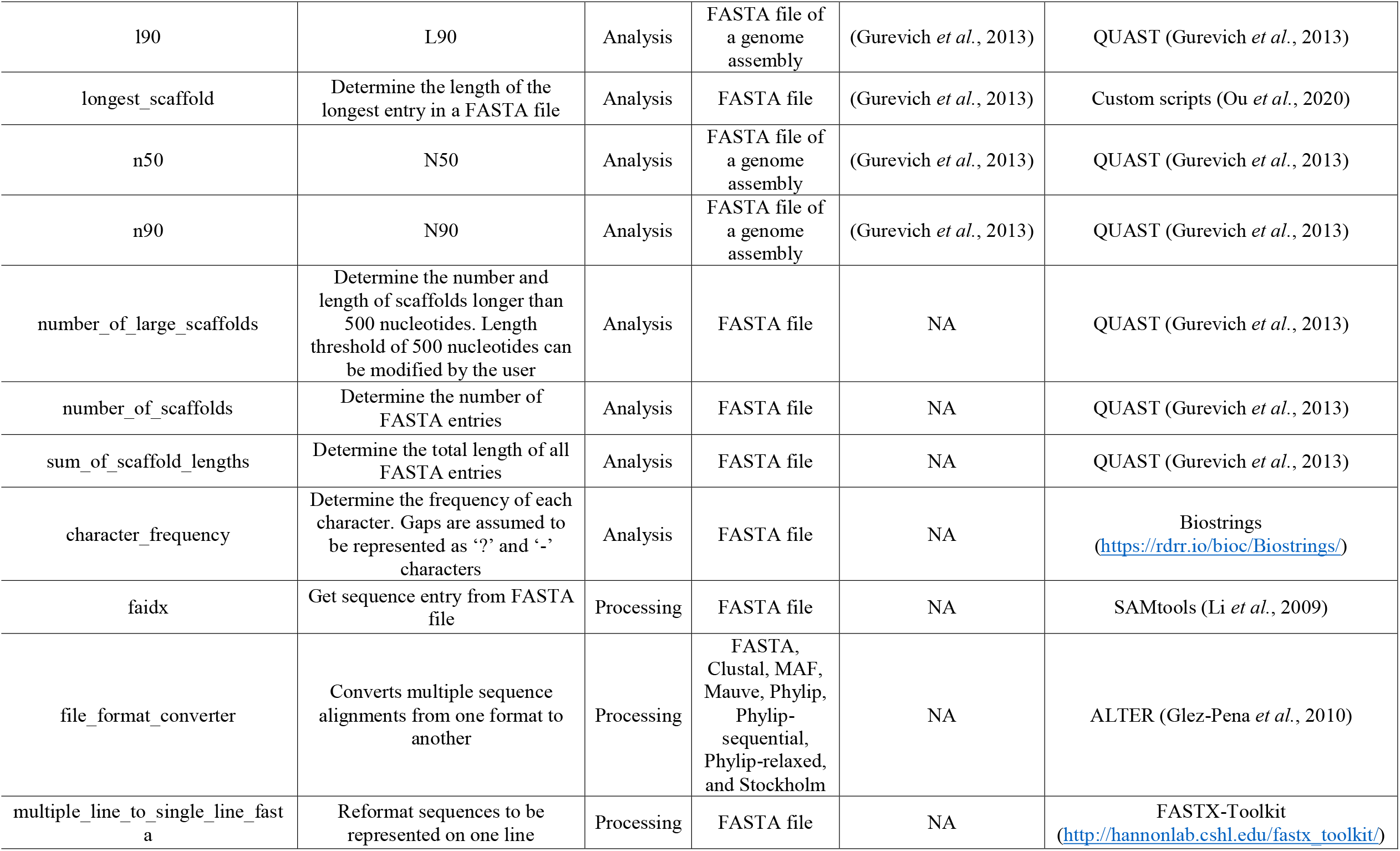
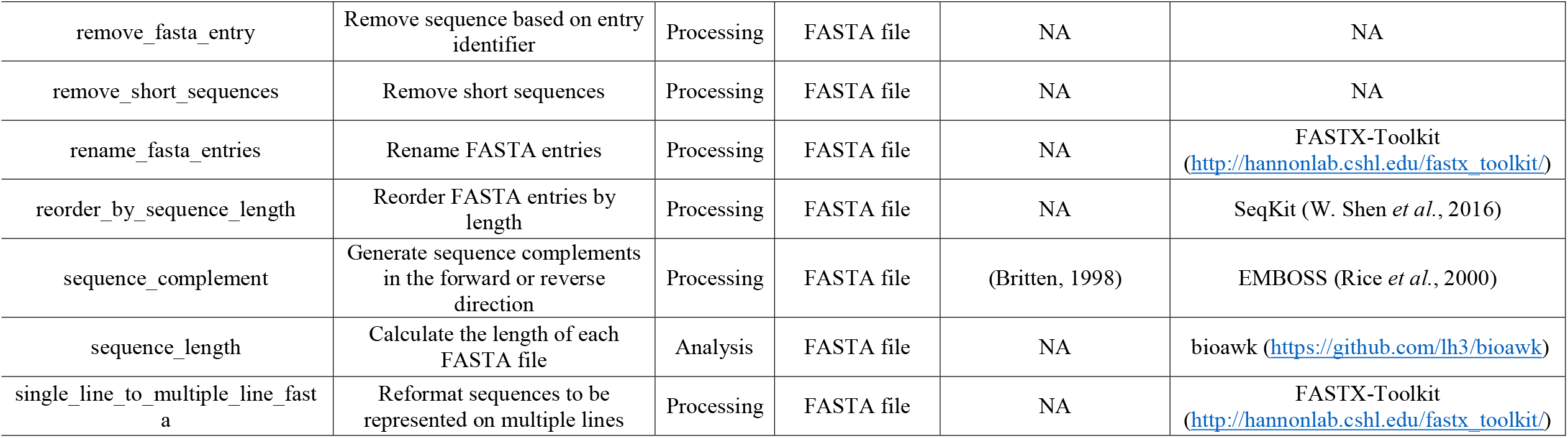
Summary of 40 functions in BioKIT.

## Materials and Methods

BioKIT is an easy-to-install command-line software that conducts diverse bioinformatic analyses in the UNIX programming environment. BioKIT is written in the Python programming language and has few dependencies, namely Biopython (Cock *et al*., 2009) and numPy (Van Der Walt *et al*., 2011).

BioKIT currently has 40 functions that process and analyze sequence files such as genome assemblies, multiple-sequence alignments, protein coding sequences, and sequencing data (Table 1). Processing functions include those that convert various file formats, subset sequence reads from FASTQ files, rename entries in FASTA files, and others. Analysis functions include those that trim sequence reads in FASTQ files according to quality and length thresholds, calculate relative synonymous codon usage, estimate codon optimization, and others. Similar to other software we have developed (Steenwyk *et al*., 2020, 2021; Steenwyk and Rokas, 2021), we plan on continuing to develop and incorporate additional functions into BioKIT to meet the needs of the research community.

Details about each function, their usage, tutorials, and other information such as how to request additional functions can be found in the online documentation (https://jlsteenwyk.com/BioKIT). To demonstrate the utility of BioKIT, we highlight four use-cases: (i) genome assembly quality assessment, (ii) summarizing properties of multiple sequence alignments, (iii) determination of relative synonymous codon usage using different genetic codes, and (iv) determination of a novel metric for estimation of gene-wise codon optimization, gene-wise relative synonymous codon usage (gw-RSCU).

### Genome assembly quality assessment

Determination of genome assembly properties is essential when evaluating assembly quality (Gurevich *et al*., 2013; Hunt *et al*., 2013). To facilitate these analyses, the *genome_assembly_metrics* function in BioKIT calculates 14 diverse properties of genome assemblies that evaluate assembly quality and characteristics including:

- assembly size: sum length of all contigs/scaffolds;
- L50 (and L90): the number of contigs/scaffolds that make up 50% (or, in the case of L90, 90%) of the total length of the genome assembly;
- N50 (and N90): the length of the contig/scaffold which, along with all contigs/scaffolds longer than or equal to that contig/scaffold, contain 50% (or, in the case of N90, 90%) the length of a particular genome assembly;
- GC content: fraction of total bases that are either G or C;
- number of scaffolds: total number of contigs/scaffolds;
- number and sum length of large scaffolds: total number and sum length of contigs/scaffolds above 500 nucleotides in length (length threshold of a “large scaffold” can be modified by the user); and
- frequency of nucleotides: fraction of occurrences for adenine (A), thymine, (T), G, and C nucleotides.

Each metric can also be called using individual functions (e.g., the *n50* function calculates the N50 of an assembly and the *number_of_large_scaffolds* function calculates the number of large scaffolds in an assembly). We anticipate the ability of BioKIT to summarize genome assembly properties will be helpful for assessing genome quality as well as in comparative studies of genome properties, such as the evolution of genome size and GC content (Walker *et al*., 2015; Shen *et al*., 2020). Other pieces of software that conduct similar analyses include QUAST, REAPR, and GenomeQC (Gurevich *et al*., 2013; Hunt *et al*., 2013; Manchanda *et al*., 2020).

### Processing and assessing the properties of multiple sequence alignments

Multiple sequence alignments—the alignment of three or more biological sequences—contain a wealth of information. To facilitate easy use and manipulation of multiple sequence alignments, BioKIT implements 16 functions that process or analyze alignments including: generating consensus sequences; generating a position-specific score matrix (which represents the frequency of observing a particular amino acid or nucleotide at a specific position); recoding an alignment using different schemes, such as the RY-nucleotide scheme for nucleotide alignments (Woese *et al*., 1991; Phillips *et al*., 2001) or the Dayhoff-6, S&R-6, and KGB-6 schemes for amino acid alignments (Hrdy *et al*., 2004; Embley *et al*., 2003; Susko and Roger, 2007; Kosiol *et al*., 2004); converting alignments among the following formats: FASTA, Clustal, MAF, Mauve, PHYLIP, PHYLIP-sequential, PHYLIP-relaxed, and Stockholm; extracting entries in FASTA files; removing entries from FASTA file; removing short sequences from a FASTA file; and others.

We highlight the *alignment_summary* function, which calculates numerous summary statistics for a multiple sequence alignment, a common step in many molecular evolutionary analyses (Plomion *et al*., 2018; Winterton *et al*., 2018). More specifically, the *alignment_summary* function calculates:

- alignment length: the total number of sites in an alignment;
- number of taxa: the total number of sequences in an alignment;
- number of parsimony-informative sites: a site in an alignment with at least two distinct nucleotides or amino acids that each occur at least twice;
- number of variable sites: a site in an alignment with at least two distinct nucleotides or amino acids;
- number of constant sites: sites with the same nucleotide or amino acid (excluding gaps); and
- the frequency of all character states: the fraction of occurrence for all nucleotides or amino acids (including gap characters represented as ‘-’ or ‘?’ in an alignment.

Like the *genome_assembly_metrics* function, each metric can be calculated individually (e.g., the *constant_sites* function calculates the number of constant sites in an alignment and the *character_frequency* function calculates the frequency of all character states). We anticipate the *alignment_summary* function will assist researchers in statistically evaluating the properties of their alignments. Other pieces of software that perform similar operations include AMAS (Borowiec, 2016) and Mesquite (Mesquite Project Team, 2014).

### Examining features of coding sequences including relative synonymous codon usage

BioKIT contains multiple functions that process or analyze protein coding sequences including translating protein coding sequences into amino acids using one of 26 genetic codes or a user-specified translation table as well as determining the GC content at the first, second, and third codon positions.

Here, we highlight the *relative_synonymous_codon_usage* function, which calculates relative synonymous codon usage, the ratio of the observed frequency of synonymous codons to an expected frequency in which all synonymous codons are used equally (Xu *et al*., 2008). In this analysis, overrepresented codons have relative synonymous codon usage values greater than one whereas underrepresented codons have relative synonymous codon usage values less than one. Relative synonymous codon usage values of one fit the neutral expectation. The *relative_synonymous_codon_usage* function can be used with one of 26 genetic codes including user-specified translation tables. The ability of BioKIT to account for diverse genetic codes makes it uniquely suitable for analyses of lineages that contain multiple genetic codes (LaBella *et al*., 2019; Krassowski *et al*., 2018). Other software that conduct similar analyses include DAMBE and GCUA (Xia, 2013; McInerney, 1998).

We also highlight the *gene_wise_relative_synonymous_codon_usage* function, which calculates a novel metric, gw-RSCU, to examine biases in codon usage among individual genes encoded in a genome. More specifically, the gw-RSCU is calculated by determining the mean or median relative synonymous codon usage value for all codons in each gene based on their genome-wide values. Thus, BioKIT calculates relative synonymous codon usage for each codon based on codon usage in an entire set of protein coding genes, individually reexamines each gene and the relative synonymous codon usage value for each codon therein, and then determines the mean or median relative synonymous codon usage value for the individual gene. The formula for the mean gw-RSCU calculation is as follows:

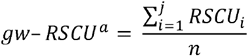

where gw-RSCU^a^ is the gene that gw-RSCU is being calculated for, RSCU_i_ is the relative synonmyouse codon usage value (calculated from all protein coding genes in a genome) for the *i*th codon of *j* codons in a gene, and *n* is the number of codons in a gene. To evaluate within-gene variation in relative synonymous codon usage, BioKIT also reports the standard deviation of relative synonymous codon usage values for each gene. Like the *relative_synonymous_codon_usage* function, gw-RSCU can be calculated using alternative genetic codes including user-specified ones. Taken together, these functions can be used individually or in tandem to investigate diverse biological phenomena, including codon usage bias (LaBella *et al*., 2019; Brandis and Hughes, 2016).

### Implementing high standards of software development

Archival instability is a concern for bioinformatic tools and threatens the reproducibility of bioinformatic research. For example, in an analysis that aimed to evaluate the “installability” of bioinformatic software, 28% of over 36,000 bioinformatic tools failed to properly install due to implementation errors (Mangul, Mosqueiro, *et al*., 2019). To ensure archival stability of BioKIT, we implemented a previously established protocol (Steenwyk and Rokas, 2021; Steenwyk *et al*., 2020, 2021) for high standards of software development and design practices. More specifically, we wrote 327 unit and integration tests that ensure faithful functionality of BioKIT and span 95.46% of the codebase. We also implemented a continuous integration pipeline, which builds, packages, installs, and tests the functionality of BioKIT across Python versions 3.6, 3.7, 3.8, and 3.9. To accommodate diverse installation workflows, we also made BioKIT freely available under the MIT license across popular platforms including GitHub (https://github.com/JLSteenwyk/BioKIT), PyPi (https://pypi.org/project/jlsteenwyk-biokit/), and the Anaconda Cloud (https://anaconda.org/jlsteenwyk/jlsteenwyk-biokit). To make BioKIT more user-friendly, we wrote online documentation, user tutorials, and instructions for requesting new features (https://jlsteenwyk.com/BioKIT). We anticipate our rigorous strategy to implement high standards of software development, coupled to our approach to facilitate easy software installation and extensive documentation, will address instabilities observed among many bioinformatic software and increase the long-term usability of BioKIT.

## Results and Discussion

### Genome assembly quality and characteristics among 901 eukaryotic genomes

To demonstrate the utility of BioKIT for the examination of genome assembly quality and characteristics, 14 diverse genome assembly metrics were determined among 901 scaffold-level haploid assemblies of eukaryotic genomes, which were obtained from NCBI, and span three major classes of animals (Mammalia; N = 350), plants (Magnoliopsida; N = 336), and fungi (Eurotiomycetes; N = 215). Genome assembly properties exhibited variation both within and between the three classes (Figure 1). For example, fungi had the smallest average genome size of 32.71 ± 7.04 Megabases (Mbs) whereas mammals had the largest average genome size of 2,645.50 ± 487.48 Mbs. Extensive variation in genome size within each class corroborates previous findings of extreme genome size variation among eukaryotes (Elliott and Gregory, 2015). Variation in GC content, a genome property that has been actively investigated for decades (Romiguier *et al*., 2010; Serres-Giardi *et al*., 2012; Galtier *et al*., 2001), was observed among the three eukaryotic classes—animals, plants, and fungi had an average GC content of 0.40 ± 0.04, 0.35 ± 0.04, and 0.49 ± 0.03, respectively. Lastly, there was wide variation in genome assembly metrics associated with continuity of assembly. For example, the average N50 values for animals, plants, and fungi were 12,287.64 ± 25,317.31 Mbs, 5,030.15 ± 19,358.58 Mbs, and 1,370.77 ± 1,552.13 Mbs, respectively. Taken together, these results demonstrate BioKIT can assist researchers in summarizing diverse genome assembly properties, which may be helpful not only for evaluating genome assembly quality, but also for studying genome evolution.

**Figure 1.**
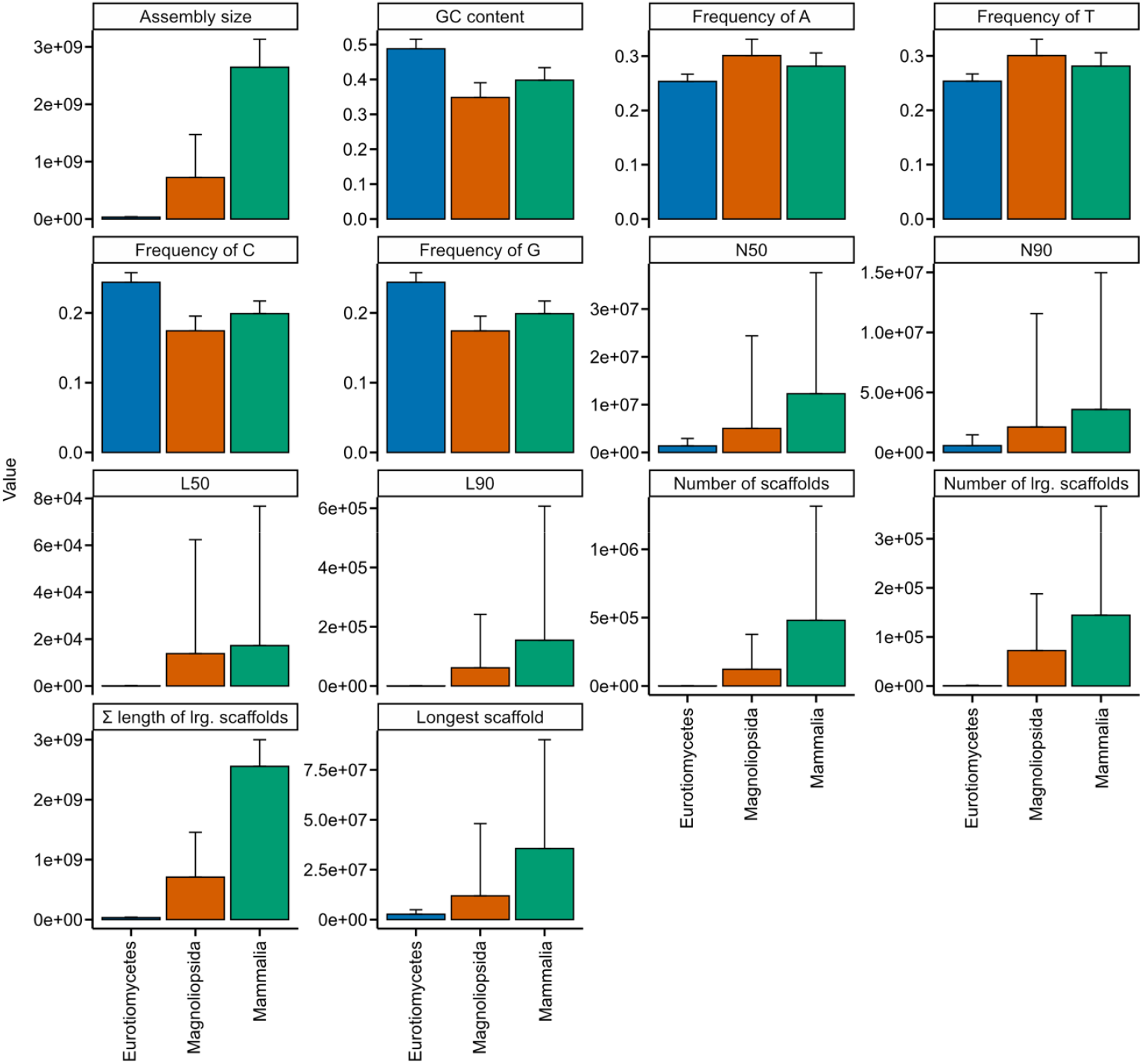
Summary of genome assembly metrics across 901 genomes from three eukaryotic classes. Nine hundred and one scaffold-level genome assemblies from three major eukaryotic classes (215 Eurotiomycetes (kingdom: Fungi), 336 Magnoliopsida (kingdom: Plantae), 350 Mammalia (kingdom: Animalia)) were obtained from NCBI and examined for diverse metrics including assembly size, GC content, frequency of A, T, C, and G, N50, N90, L50, L90, number of scaffolds, number of large scaffolds (defined as being greater than 500 nucleotides, which can be modified by the user), sum length of large scaffolds, and longest scaffold in the assembly. Bar plots represent the mean for each taxonomic class. Error bars represent the standard deviation of values. This figure was made using ggplot2 (Wickham, 2009) and ggpubfigs (https://github.com/JLSteenwyk/ggpubfigs).

### Properties of multiple sequence alignment from 10 phylogenomic studies

To demonstrate the utility of BioKIT in calculating summary statistics for multiple sequence alignments, we calculated six properties across 10 previously published phylogenomic data matrices of amino acid sequences (Borowiec *et al*., 2015; Chen *et al*., 2015; Misof *et al*., 2014; Nagy *et al*., 2014; Shen *et al*., 2018; X.-X. Shen, Zhou, *et al*., 2016; Steenwyk *et al*., 2019; Struck *et al*., 2015; Whelan *et al*., 2015; Yang *et al*., 2015) (Figure 2). Phylogenomic data matrices varied in the number of taxa (mean = 109.50 ± 87.26; median = 94; max = 343; min = 36). Alignment length is associated with greater phylogenetic accuracy and bipartition support (X.-X. Shen, Salichos, *et al*., 2016); however, recent analyses suggest that in some instances shorter alignments that contain a wealth of informative sites (such as parsimony-informative sites) harbor robust phylogenetic signal (Steenwyk *et al*., 2020). Interestingly, the longest observed alignment (1,806,035 sites; *Chen, Vertebrates* in Figure 2) (Chen *et al*., 2015) contained the highest number of constant sites (N = 610,994), which are phylogenetically uninformative, as well as the highest number of variable sites (N = 1,195,041), which are phylogenetically informative (X.-X. Shen, Salichos, *et al*., 2016). In contrast to the multiple sequence alignment of vertebrate sequences, the second longest alignment of budding yeast sequences (1,162,805 sites; *Shen, 332 Yeast* in Figure 2) has few constant sites (N = 2,761) and many parsimony-informative (N = 1,152,145) and variable sites (N = 1,160,044). This observation may be driven in part by the rapid rate of budding yeast evolution compared to animals (Shen *et al*., 2018). These results demonstrate BioKIT is useful in summarizing multiple sequence alignments.

**Figure 2.**
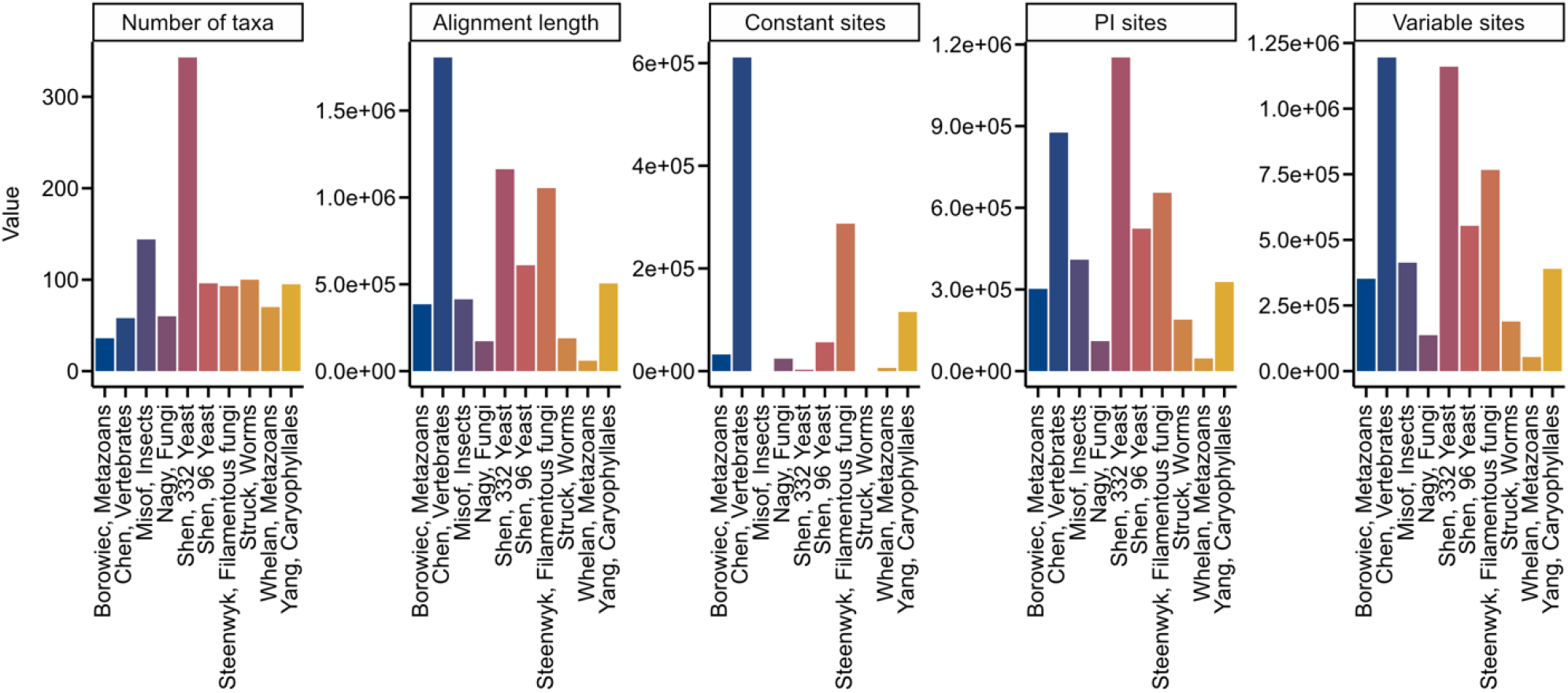
Summary metrics among multiple sequence alignments from phylogenomic studies. Multiple sequence alignments of amino acid sequences from ten phylogenomic data matrices (Borowiec *et al*., 2015; Chen *et al*., 2015; Misof *et al*., 2014; Nagy *et al*., 2014; Shen *et al*., 2018; X.-X. Shen, Zhou, *et al*., 2016; Steenwyk *et al*., 2019; Struck *et al*., 2015; Whelan *et al*., 2015; Yang *et al*., 2015) were examined for five metrics: number of taxa, alignment length, number of constant sites, number of parsimony-informative sites, and number of variable sites. The x-axis depicts the last name of the first author of the phylogenomic study followed by a description of the organisms that were under study. The abbreviation PI represents parsimony-informative sites. Although excluded here for simplicity and clarity, BioKIT also determines character state frequency (nucleotide or amino acid) when summarizing alignment metrics. This figure was made using ggplot2 (Wickham, 2009) and ggpubfigs (https://github.com/JLSteenwyk/ggpubfigs).

### Relative synonymous codon usage in 107 budding yeast and filamentous fungi

To demonstrate the utility of BioKIT in analyzing protein coding sequences, we calculated the relative synonymous codon usage of all codons in the protein coding sequences of 103 Eurotiomycetes (filamentous fungi) and 68 Saccharomycetes (budding yeasts) genomes obtained from the RefSeq database of NCBI (Figure 3). This example also demonstrates the flexibility of BioKIT to account for non-standard genetic codes, which are observed among some budding yeasts that use the CUG codon to encode a serine or alanine rather than a leucine (Krassowski *et al*., 2018). Hierarchical clustering of relative synonymous codon usage values per codon (columns in Figure 3) revealed similar patterns across groups of codons. For example, CUA, AUA, and GUA—three of the four codons that end in UA—were underrepresented in all fungi. Hierarchical clustering of relative synonymous codon usage values per species (rows in Figure 3) revealed filamentous fungi and budding yeasts often clustered separately. For example, UGA, GUG, AAC, UAC, AAG, UUC, UCC, ACC, GCC, CGC, CUG, AUC, GUC, CUC, and GGC are more often overrepresented among filamentous fungi in comparison to budding yeasts; in contrast, UUG, GUU, CCA, and GGU are more often overrepresented among budding yeasts in comparison to filamentous fungi. Variation within each lineage was also observed; for example, UUA was underrepresented in most, but not all, budding yeasts.

**Figure 3.**
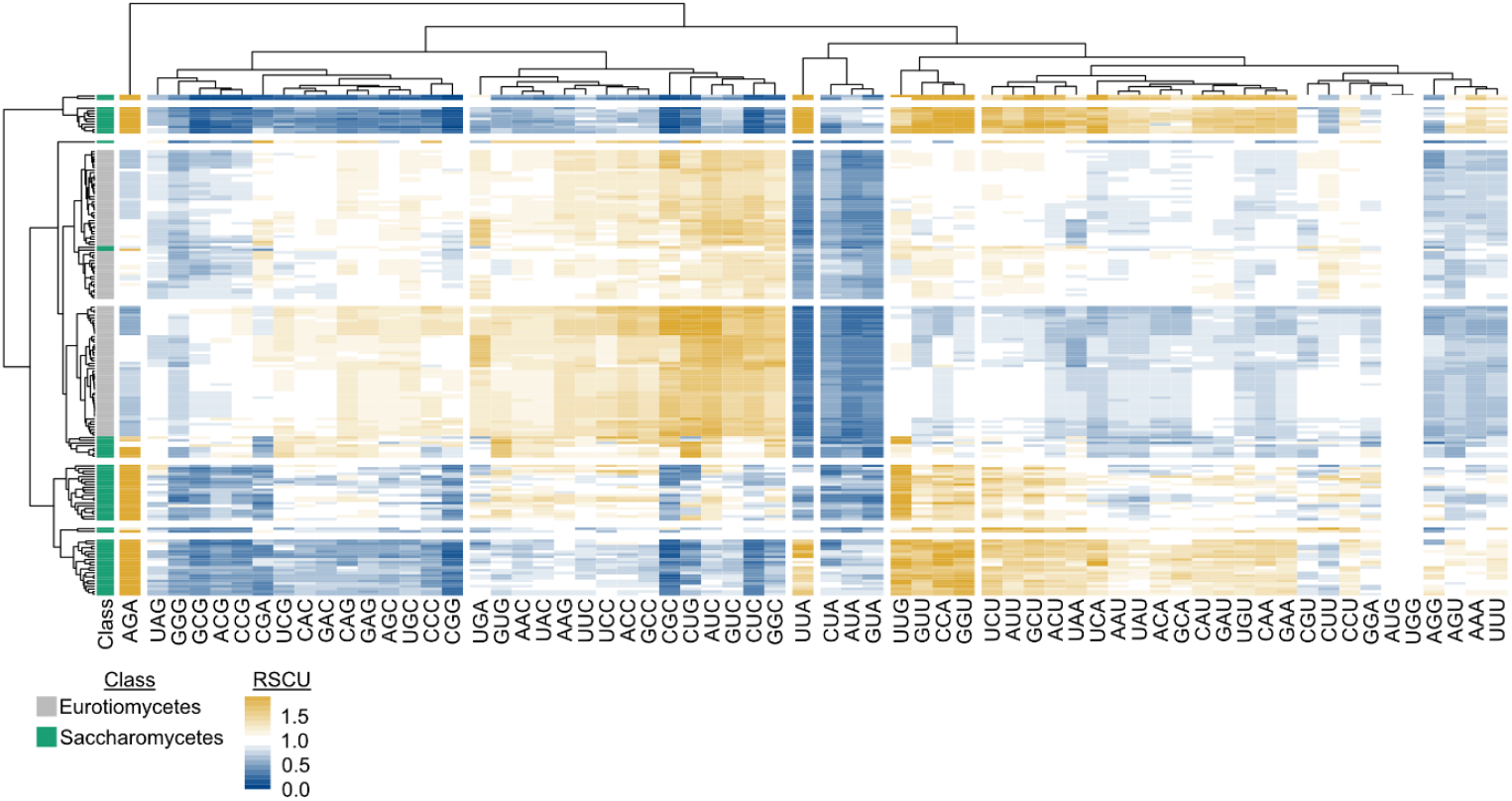
Relative synonymous codon usage across 171 fungal genomes. Relative synonymous codon usage (RSCU) was calculated from the coding sequences of 103 Eurotiomycetes (filamentous fungi) and 68 Saccharomycetes (budding yeasts) genomes obtained from NCBI. Hierarchical clustering was conducted across the fungal species (rows) and codons (columns). Eight groups of clustered rows were identified; seven groups of clustered columns were identified. Broad differences were observed in the RSCU values of Eurotiomycetes and Saccharomycetes genomes. For example, Saccharomycetes tended to have higher RSCU values for the AGA codon, whereas Eurotiomycetes tended to have higher RSCU values for the CUG codon. To account for the use of an alternative genetic code in budding yeast genomes from the CUG-Ser1 and CUG-Ser2 lineages (Krassowski *et al*., 2018), the alternative yeast nuclear code—which is one of 26 alternative genetic codes incorporated into BioKIT—was used during RSCU determination. User’s may also provide their own genetic code if it is unavailable in BioKIT. Overrepresented codons (RSCU>1) are depicted in a gold gradient; underrepresented codons (RSCU<1) are depicted in a blue gradient. RSCU values greater than 2 are depicted with the maximum gold color. Eurotiomycetes are depicted in grey; Saccharomycetes are depicted in green. This figure was made using pheatmap (Kolde, 2012).

### Patterns of gene-wise codon usage bias can be used to assess codon optimization and predict steady-state gene expression levels

To evaluate the utility of BioKIT in examining gene-wise codon usage biases, we calculated the mean and median gw-RSCU value, a novel metric introduced in the present manuscript, for individual protein coding genes in the genome of *S. cerevisiae* (Figure 4A). Mean and median gw-RSCU values were often, but not always, similar—the average absolute difference between mean and median gw-RSCU is 0.05 ± 0.04. In *S. cerevisiae*, as well as other organisms, genes encoding ribosomal components and histones are known to be codon optimized and highly expressed (Hershberg and Petrov, 2009; LaBella *et al*., 2021; Sharp *et al*., 1986). Therefore, we hypothesized that genes with high gw-RSCU values will have functions related to ribosomes or histones because patterns of gene-wise codon usage bias may be indicative of codon optimization. Supporting this hypothesis, examination of the 10 genes with the highest mean gw-RSCU revealed five genes with ribosome-related functions [RPL41B (YDL133C-A), mean gw-RSCU: 1.60; RPL41A (YDL184C), mean gw-RSCU: 1.58; RPS14A (YCR031C), mean gw-RSCU: 1.44; RPS9B (YBR189W), mean gw-RSCU: 1.43; and RPL18A (YOL120C), mean gw-RSCU: 1.43] and four genes with histone-related functions [HHF1 (YBR009C), mean gw-RSCU: 1.45; HTA2 (YBL003C), mean gw-RSCU: 1.44; HHF2 (YNL030W), mean gw-RSCU: 1.43; and HTA1 (YDR225W), mean gw-RSCU: 1.43]. Examination of the 10 most optimized genes according to median gw-RSCU revealed similar observations wherein nine genes had ribosome-related functions [RPS14A (YCR031C), median gw-RSCU: 1.48; RPS12 (YOR369C), median gw-RSCU: 1.40; RPS30B (YOR182C), median gw-RSCU: 1.40; RPP2A (YOL039W), median gw-RSCU: 1.40; RPL18A (YOL120C), median gw-RSCU; RPS3 (YNL178W), median gw-RSCU: 1.40; RPL13B (YMR142C), median gw-RSCU: 1.40; RPP0 (YLR340W), median gw-RSCU: 1.40; and RPS0B (YLR048W), median gw-RSCU: 1.40]. More broadly, genes associated with the 60S and 40S ribosomal units (gold color in Figure 4A) tended to have high gw-RSCU values. These results suggest gw-RSCU values may be useful for estimating codon optimization. To further explore the relationship between gw-RSCU and codon optimization, we compared gw-RSCU values to the values of the tRNA adaptation index, a measure of codon optimization (Sabi and Tuller, 2014), in *S. cerevisiae* as well as in steady state gene expression data from *Saccharomyces mikatae* (LaBella *et al*., 2019). In *S. cerevisiae*, strong correlation was observed between mean gw-RSCU and tRNA adaptation index values (Figure 4B) and a less robust, but still significant, correlation was observed between median gw-RSCU and tRNA adaptation index values (Figure 4C). Examination of gw-RSCU and gene expression data from *S. mikatae* revealed a robust correlation (Figure 4E and 4F) suggesting gw-RSCU, and in particular the mean gw-RSCU, can serve as a measure of gene-wise codon optimization.

**Figure 4.**
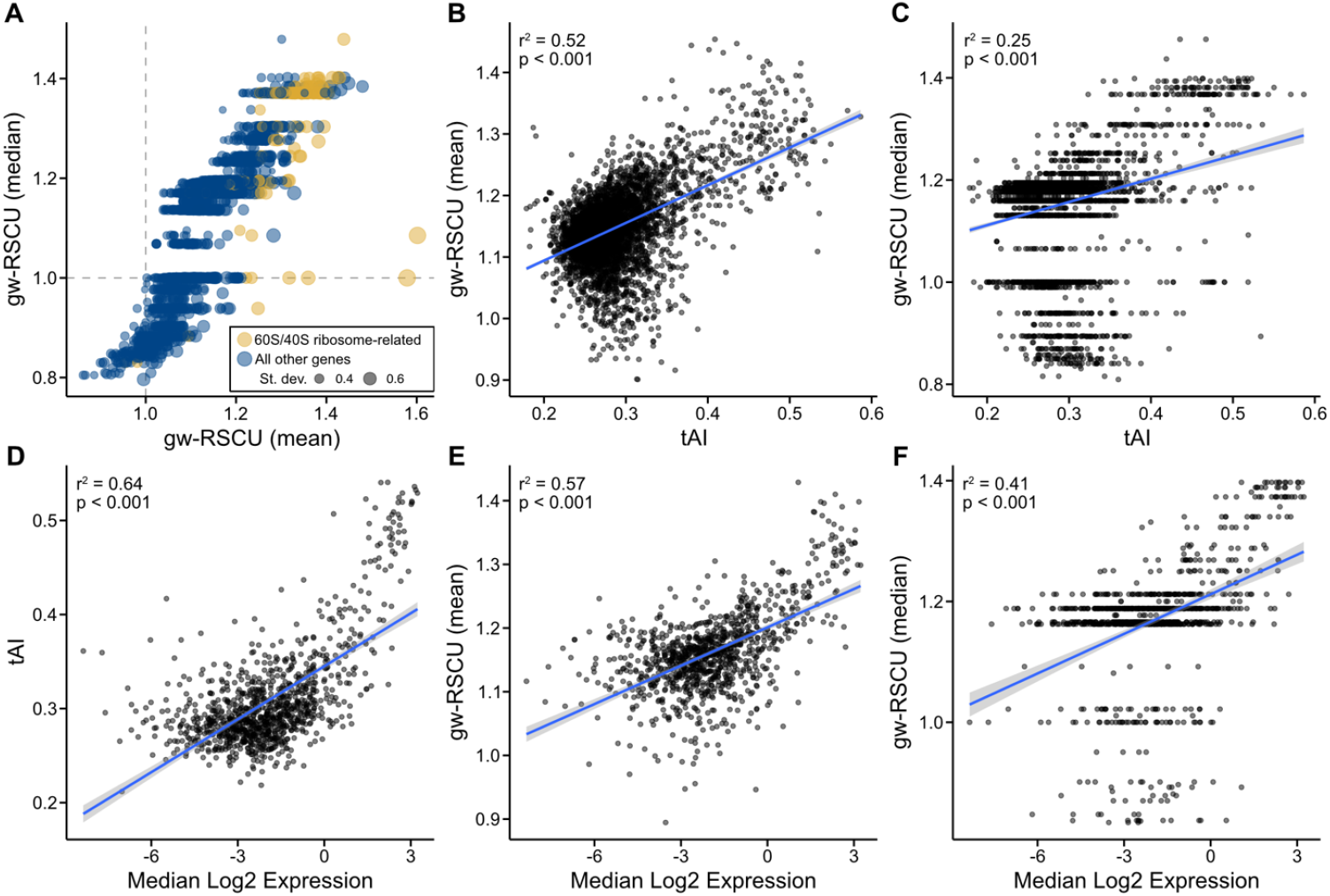
Mean gene-wise relative synonymous codon usage accurately estimates codon optimization. (A) Gene-wise relative synonymous codon usage (gw-RSCU), the mean (x-axis) or median (y-axis) relative synonymous codon usage value per gene (based on RSCU values calculated from the entire set of protein coding genes), was calculated from the coding sequences of the model budding yeast *Saccharomyces cerevisiae*. (B, C) In *S. cerevisiae*, a significant correlation was observed between tRNA adaptation index (tAI), a well-known measure of codon optimization (Sabi and Tuller, 2014), and mean as well as median gw-RSCU (r^2^ = 0.52, p < 0.001 and r^2^ = 0.25, p < 0.001, respectively; Pearson’s Correlation Coefficient). (D) Using previously published data, a correlation is observed between median log2 gene expression and tAI in *Saccharomyces mikatae* (LaBella *et al*., 2019), which is evidence of tAI values being indicative of codon optimization. Comparison of mean and median gw-RSCU (E and F, respectively) and median log2 gene expression revealed similarly strong correlations (r^2^ = 0.57, p < 0.001 and r^2^ = 0.41, p < 0.001, respectively; Pearson’s Correlation Coefficient). Of note, mean gw-RSCU had a strong correlation to gene expression than median gw-RSCU. Each gene is represented by a dot. In panel A, the size of each dot represents the standard deviation of RSCU values observed in the gene and the color of each dot represents if the protein encoded by the gene has functions related to the 60S and 40S ribosomal subunits (gold) or a different function (blue).

## Conclusion

BioKIT is a multi-purpose toolkit that has diverse applications for bioinformatics research. The utilities implemented in BioKIT aim to facilitate the execution of seamless bioinformatic workflows that handle diverse sequence file types. Implementation of state-of-the-art software development and design principles in BioKIT help ensure faithful function and archival stability. BioKIT will be helpful for bioinformaticians with varying levels of expertise and biologists from diverse disciplines including molecular biology.

## Data Availability

BioKIT is freely available under the MIT license from GitHub (https://github.com/JLSteenwyk/BioKIT), PyPi (https://pypi.org/project/jlsteenwyk-biokit/), and the Anaconda Cloud (https://anaconda.org/jlsteenwyk/jlsteenwyk-biokit). Documentation, user tutorials, and instructions for requesting new features are available online (https://jlsteenwyk.com/BioKIT).

## Acknowledgements

We thank the Rokas lab for helpful discussion and feedback. We also thank the bioinformatic community for providing suggestions of functions to implement in BioKIT. J.L.S. and A.R. were funded by the Howard Hughes Medical Institute through the James H. Gilliam Fellowships for Advanced Study program. Research in A.R.’s lab is supported by grants from the National Science Foundation (DEB-1442113 and DEB-2110404), the National Institutes of Health/National Institute of Allergy and Infectious Diseases (R56 AI146096), and the Burroughs Wellcome Fund.

## Conflict of Interest

A.R. is a scientific consultant for LifeMine Therapeutics, Inc.

## Notes

https://github.com/JLSteenwyk/BioKIT

### Competing Interest Statement

Antonis Rokas is a scientific consultant for LifeMine Therapeutics, Inc.

### Summary of Updates

Some of the links to the software provided in v1 were incorrect. The revised version has fixed these errors.

